## Supplementary Materials for "Mutation rate dynamics reflect ecological change in an emerging zoonotic pathogen"

**This PDF file includes:**

Figures S1 to S13, Tables S1, S3, S4, S6-7, S10-11, S15, Legends for Tables S2, S5, S8-9, S12-14, S16, SI References

**Other supplementary materials for this manuscript include the following:**

Tables S2, S5, S8-9, S12-14, S16

### Supplementary Figures

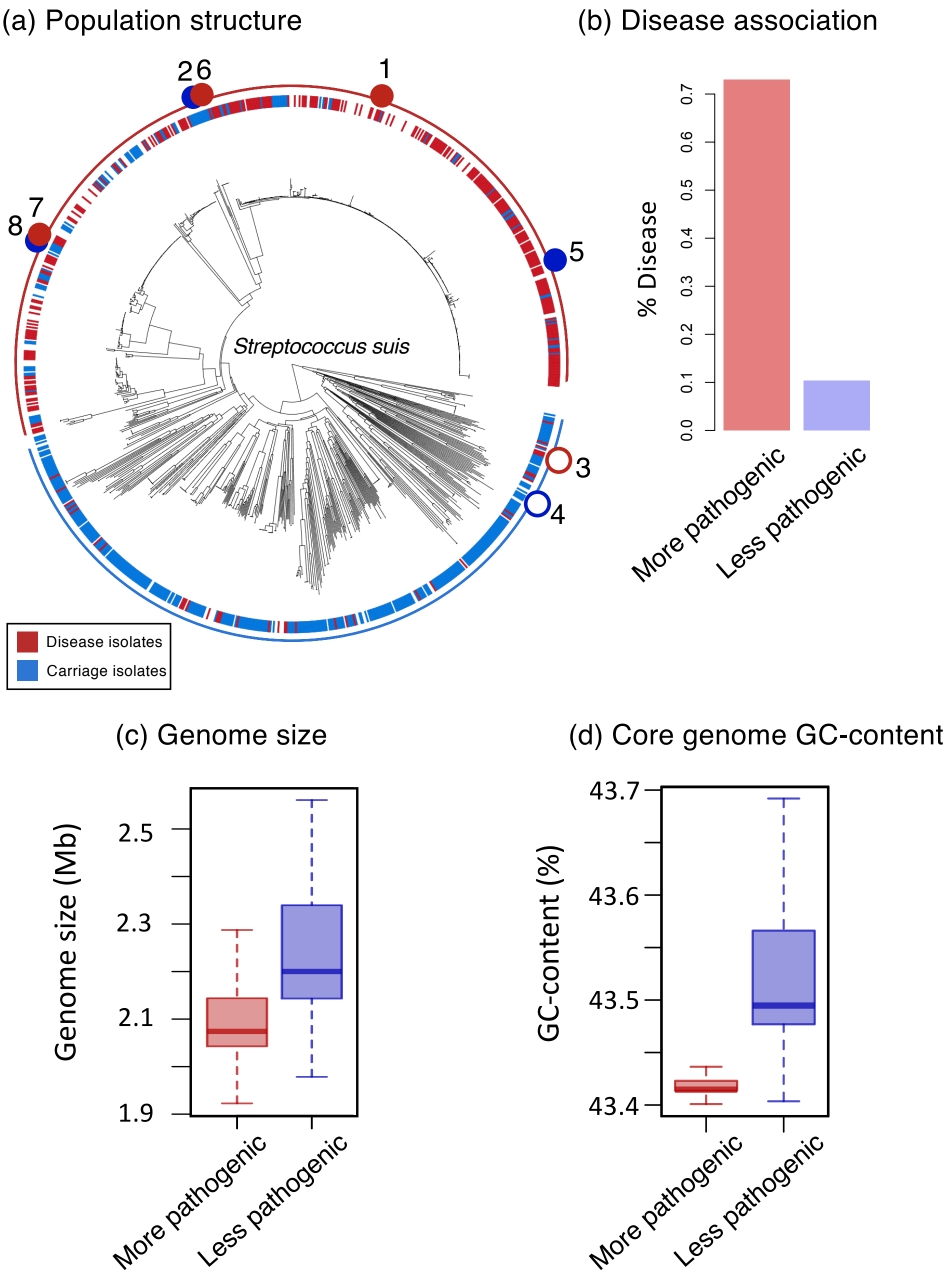


**Fig. S1. Population structure of a global collection of *Streptococcus suis* isolates.** (a) A core genome phylogeny of 962 isolates of *S. suis*[1]. Individual disease (red) and carriage (blue) isolates are indicated in the inner strip. The more pathogenic clade is indicated by a red outer ring, and the less pathogenic clade by a blue outer ring. The locations of each strain in our two MA experiments are indicated on this strip (Table 1). (b) The proportion of isolates in each clade that are associated with disease (excluding isolates for which disease-association is unknown). (c) A box plot of genome sizes of isolates from each clade. (d) A box plot of the core genome GC-content of isolates from each clade.

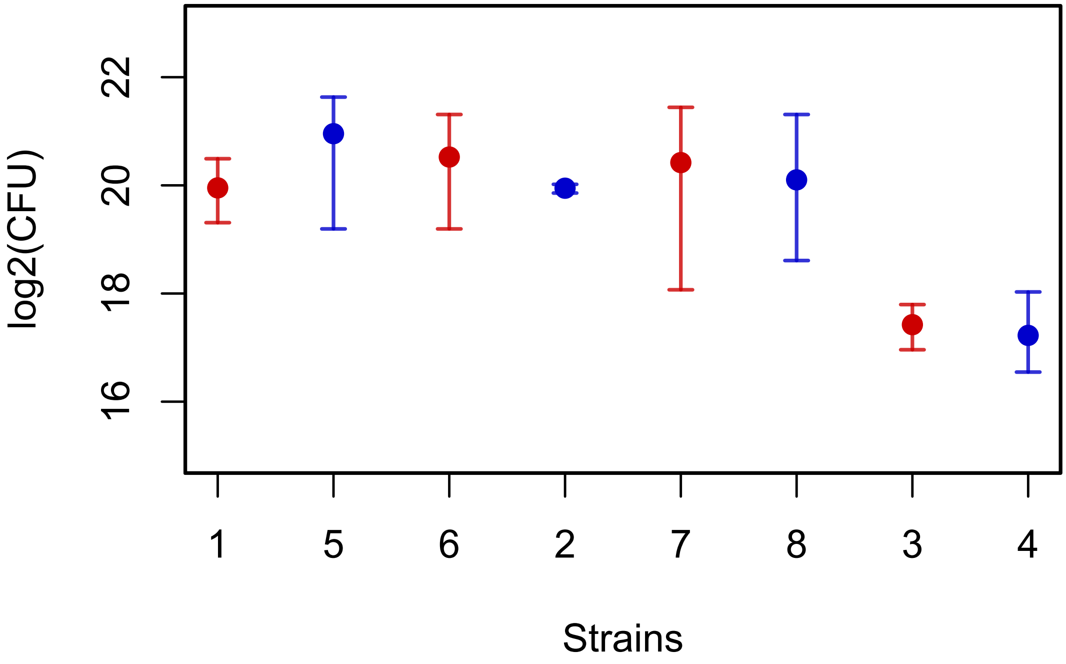


**Fig. S2. Estimates of number of generations per day based on colony counts for each ancestral strain.** Points represent mean estimates of the number of colony forming units (CFU) present after 24 hours of growth from a single CFU for each ancestral strain, on a log2 scale. Bars show the range of values returned across biological replicates (at least 3 biological replicates of each strain). Disease strains are shown in red and carriage strains in blue. Log2(CFU) after 24 hours of growth gives an estimate of the number of generations over the that period. We find no evidence of a difference in generation time between disease and carriage strains, but the two strains from the less pathogenic clade (strains 3 and 4) have a longer generation time than the six strains from the more pathogenic clade (strains 1, 2, 5, 6, 7 and 8).

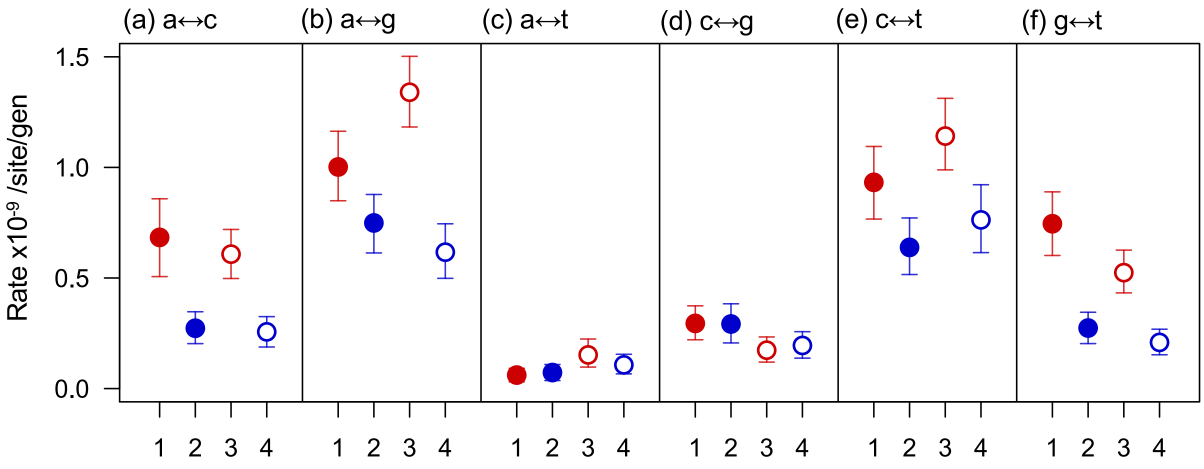


**Fig. S3. Comparison of mutation rates for different types of single nucleotide variants in the 200-day MA experiment.** Points represent mean values across 50 replicate lines, and bars represent 95% confidence intervals estimated from bootstrapping across lines. Numbers relate to Table 1; disease strains are shown in red and carriage in blue, strains from the more pathogenic clade are shown as filled shapes and strains from the less pathogenic clade as unfilled shapes.

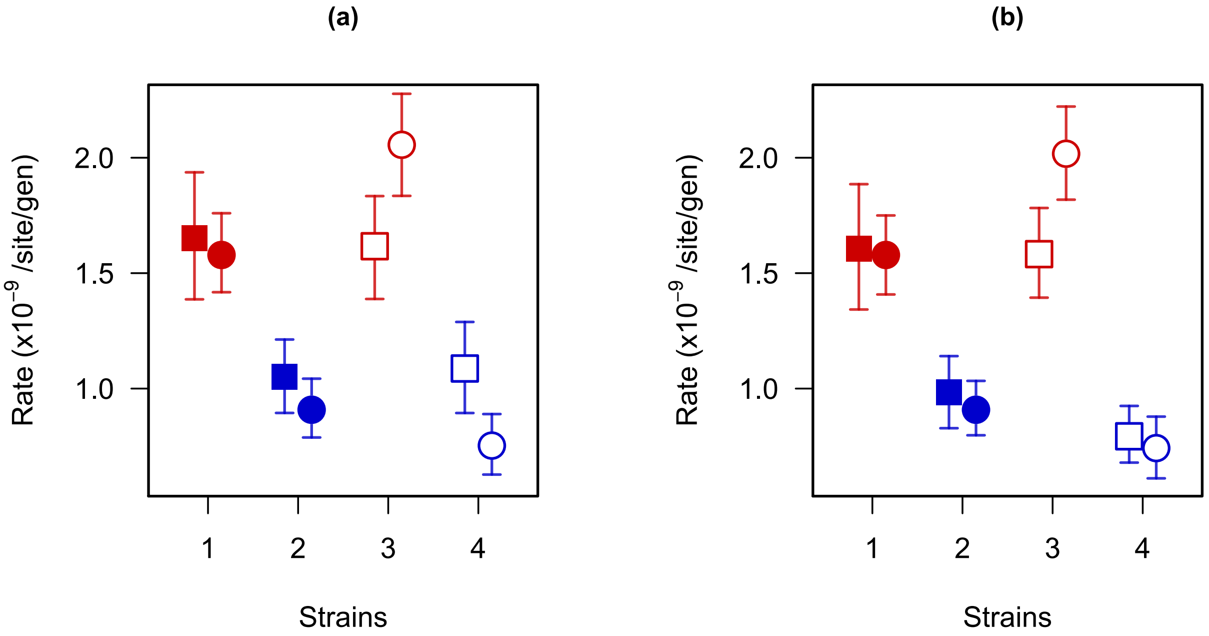


**Fig. S4. Comparison of mutation rates across core and accessory genes in the 200-day MA experiment, with (a) and without clustered mutations (b).** Estimates of rates of single-base mutation rates across accessory (squares) and core (circles) genes. Points represent mean values across 50 replicate lines, and bars represent 95% confidence intervals estimated from bootstrapping across lines. Numbers relate to Table 1; disease strains are shown in red and carriage in blue, strains from the more pathogenic clade are shown as filled shapes and strains from the less pathogenic clade as unfilled shapes.

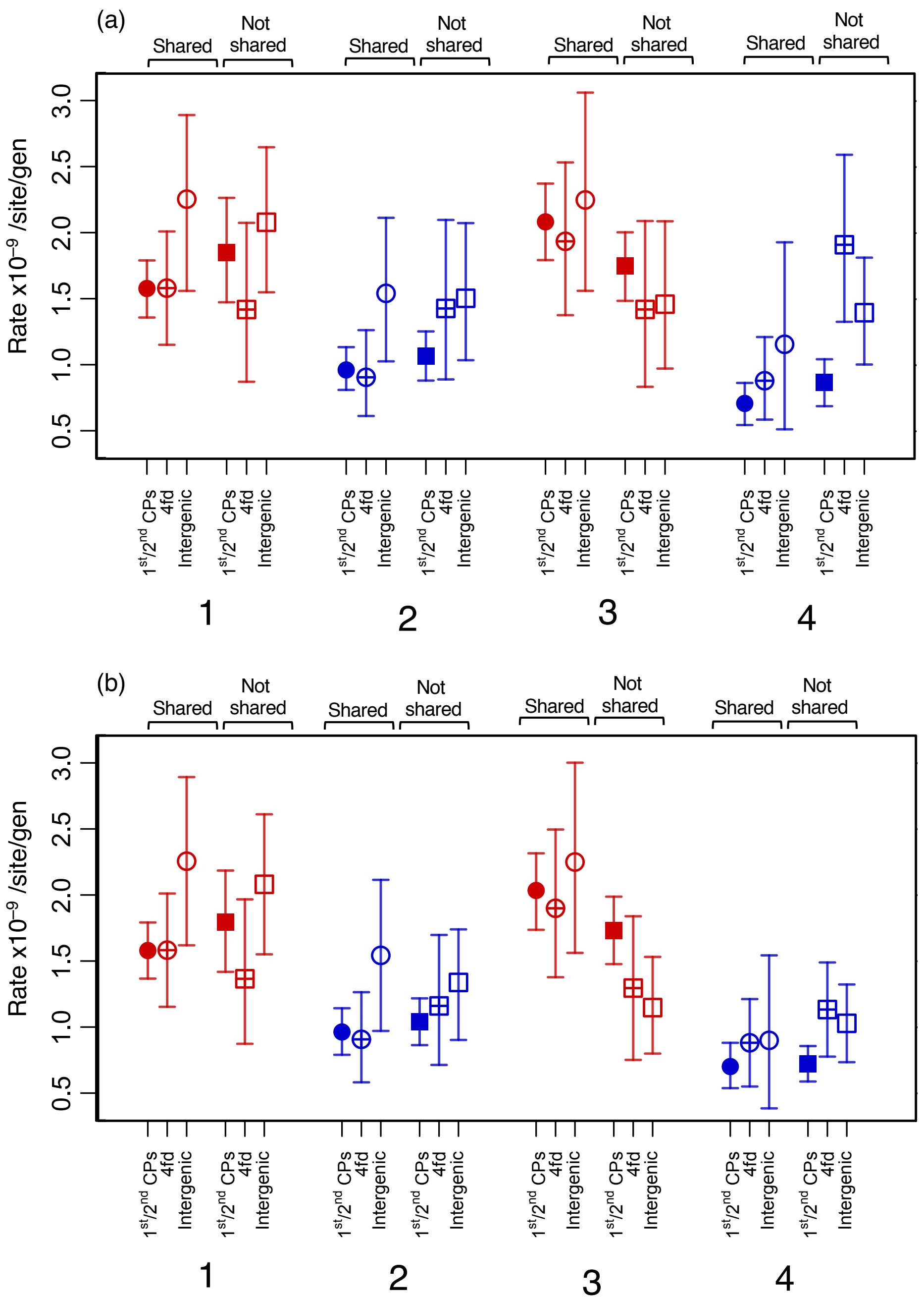


**Fig. S5. Comparison of mutation rates in the 200-day MA experiment across different categories of sites, with (a) and without clustered mutations (b).** Both figures show rates for 1st/2nd codon positions, four-fold degenerate sites, and intergenic sites, for both regions that are shared across all four strains and regions that are not shared. Shared intergenic sites were defined as intergenic regions between genes that were syntenic across all four strains. Estimates based only on genes that are shared across all four strains (core) are shown as circles, and estimates based only on genes that are not shared across all four strains (accessory) are shown as squares. Disease strains are shown in red and carriage strains in blue. All points represent mean values across 50 replicate lines, and bars represent 95% confidence intervals estimated from bootstrapping across lines.

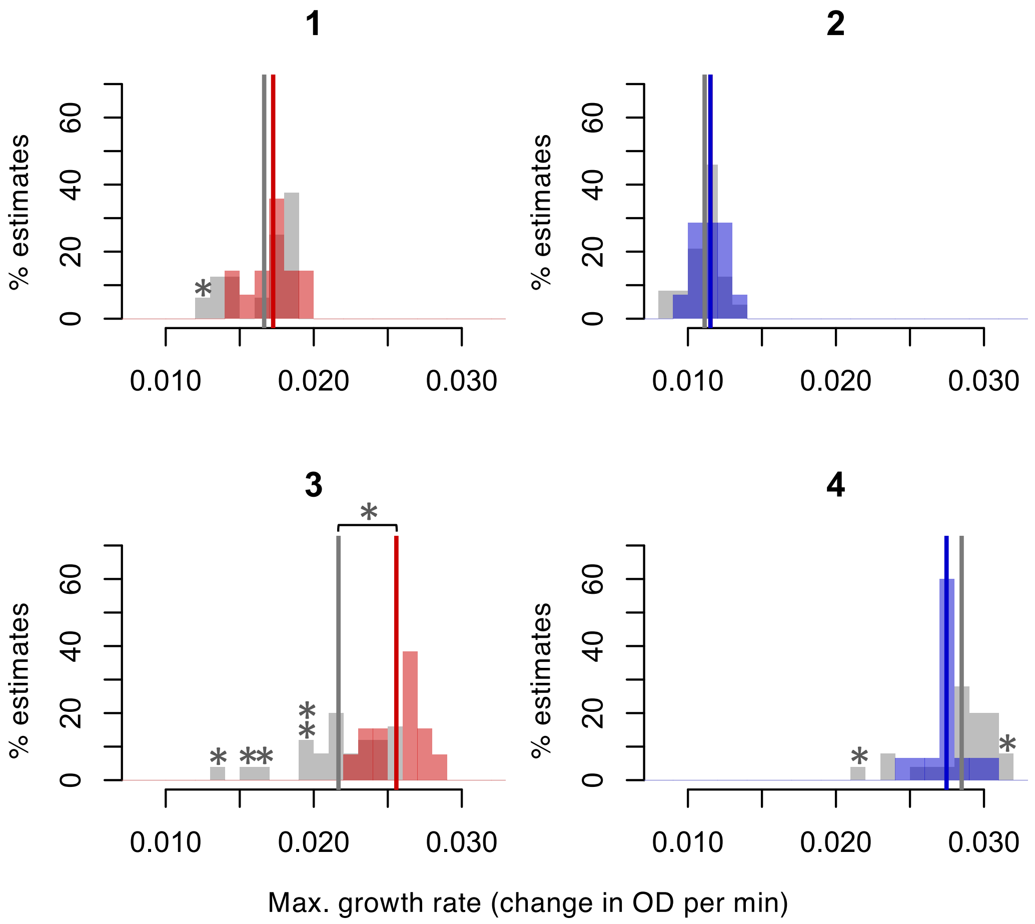


**Fig. S6. Comparison of the maximum growth rate estimates for evolved and ancestral lines of the four strains in the 200-day MA experiments.** Histograms of maximum growth rates (change in optical density (OD) per min) for evolved lines (mean values across 3 biological replicates; shown in grey) and repeated measurements of the ancestral line (red for disease and blue for carriage). Estimates were attempted for a random sample of 25 evolved lines for each strain, however insufficient overnight growth meant that we could not obtain accurate maximum growth rate estimates for 9/25 evolved lines of strain 1 and 1/25 evolved lines of strain 2. Estimates for these lines are therefore not shown. Vertical lines represent mean values across repeated measurements of the ancestral strain (red/blue), and across evolved lines (grey). For strain 3 there is evidence of a net decline in maximum growth rate in the evolved lines (Welch's t-test, $p = 2.9 \times 10^{-5}$, indicated by * above brackets). For the three other strains there is no evidence of a net change in the maximum growth rate. Estimates of maximum growth rate for 7 individual evolved lines were significantly lower than estimates for the ancestral strain, and 1 significantly higher (Welch's t-test, $p < 0.05$ after Bonferroni correction for multiple testing, indicated by * above bars).

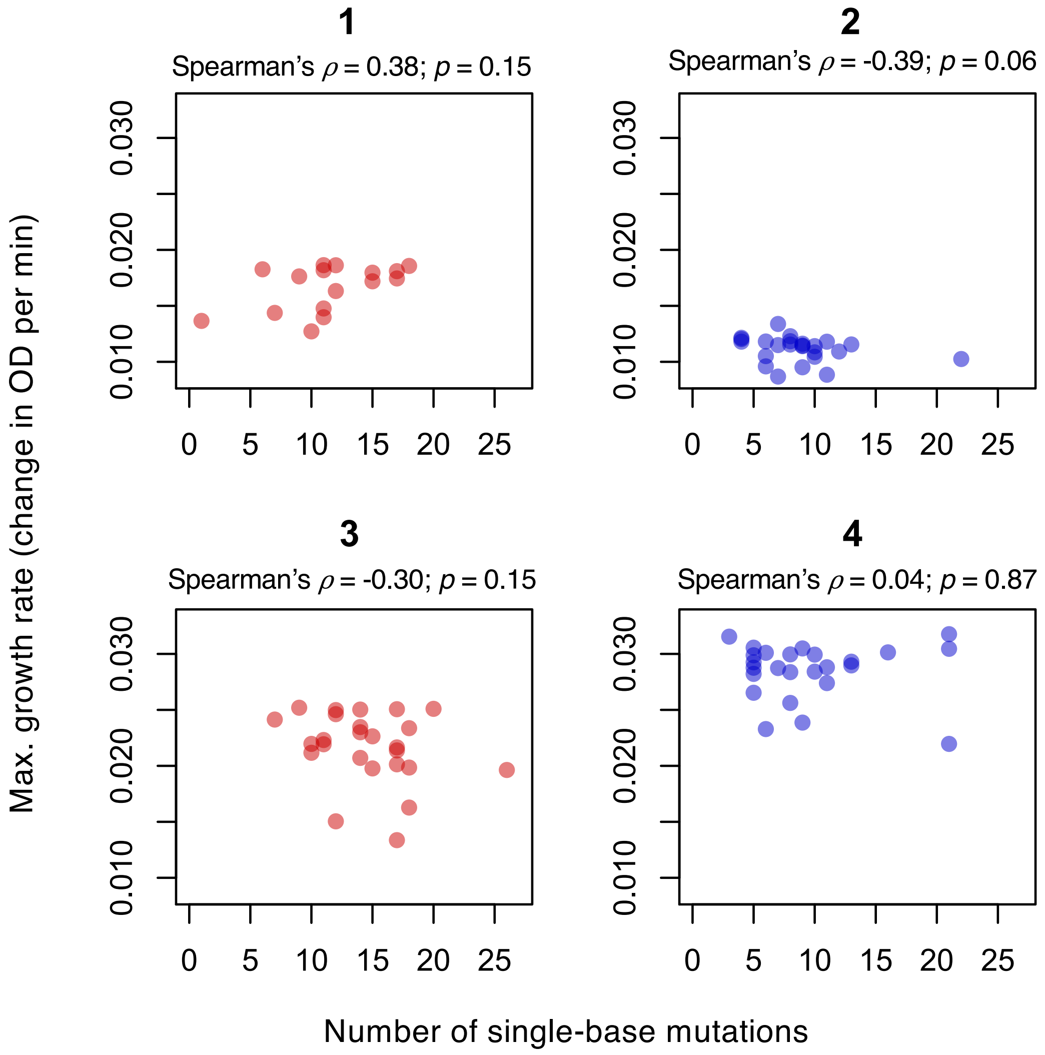


**Fig. S7. Relationship between maximum growth rate and numbers of single-base mutations accumulated in the evolved lines of the four strains in the 200-day MA experiments.** Maximum growth rate (change in optical density (OD) per min) plotted against the number of single-base substitutions accumulated over the course of the experiment for evolved lines of each strain. Growth rate estimates were attempted for a random sample of 25 evolved lines of each strain, however insufficient overnight growth meant that we could not obtain accurate maximum growth rate estimates for 9/25 lines of strain 1 and 1/25 lines of strain 2. The estimates of maximum growth rates shown are mean values across 3 biological replicates. Spearman's $\rho$ and p-values for correlation tests are reported for each strain. There is no evidence of a significant correlation between maximum growth rate and the number of mutations accumulated for any of our four strains. However, the 9 lines of strain 1 that had insufficient overnight growth had more single-base mutations (average of 15.4) than 16 lines that had sufficient overnight growth (average of 11.4) (Welch's t-test, $p = 0.04$). Similarly, the line of strain 2 than had insufficient overnight growth had a high number of single-base mutations (13) compared to the other lines of this strain included in the experiment.

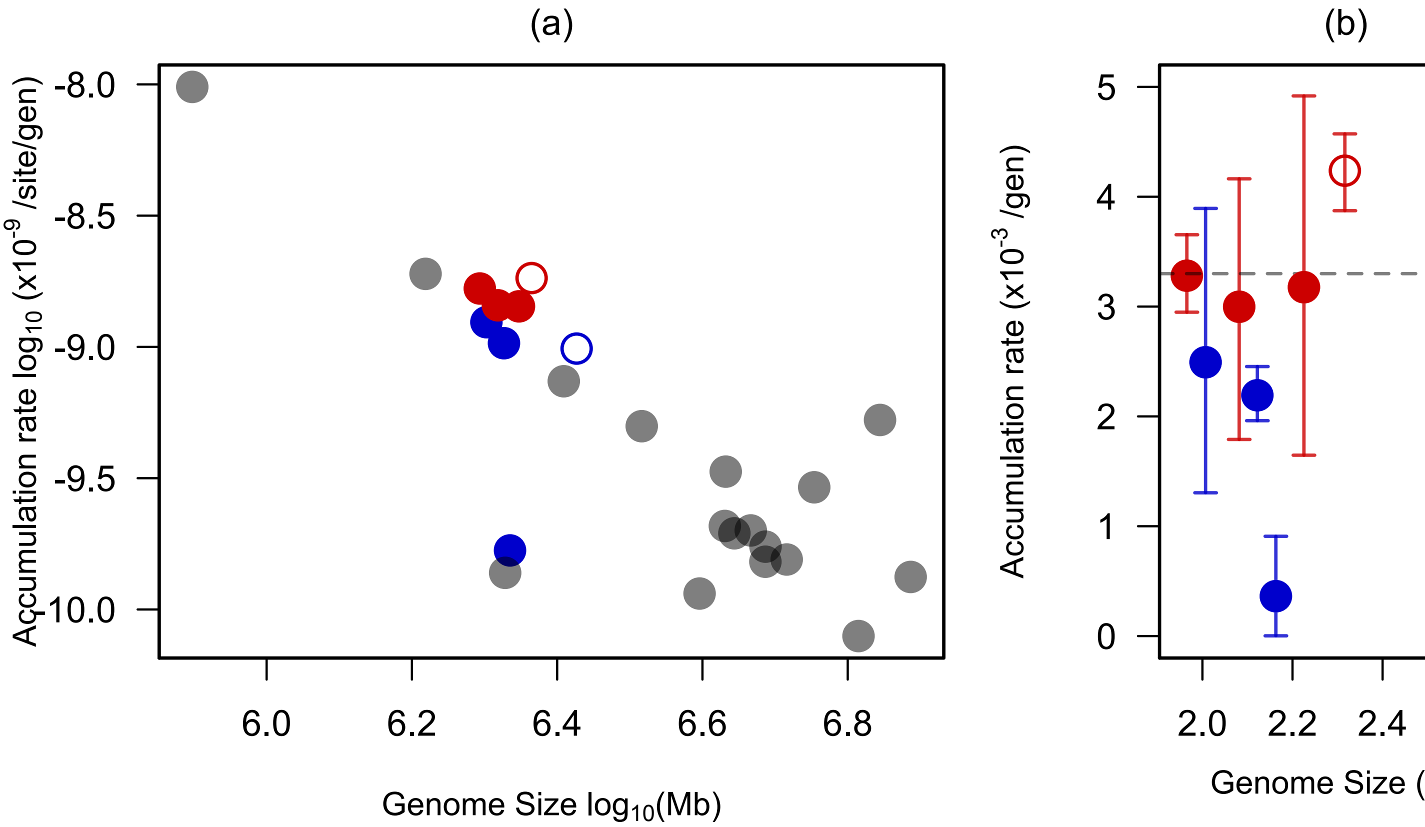


**Fig. S8. (a) Mutation rate estimates for *S. suis* in the context of those of other species, and (b) a comparison of mutation rate estimates per genome.** (a) Mutation rate estimates against genome size, for *S. suis*, *Mesoplasma florum, Helicobacter pylori, Thermus thermophilus, Staphylococcus epidermidis, Deinococcus radiodurans, Vibrio cholerae, Vibrio fischeri, Bacillus subtilis, Mycobacterium tuberculosis, Escherichia coli, Salmonella typhimurium, Salmonella enterica, Teredinibacter turnerae, Agrobacterium tumefaciens, Pseudomonas aeruginosa, Mycobacterium smegmatis* and *Burkholderia cenocepacia*[2,3]. *S.* s*uis* disease isolates are shown in red and carriage isolates in blue, with closed circles representing isolates from the more pathogenic clade and open circles isolates from a less pathogenic clade, other species are shown in grey. In (b) the dashed line indicates the value of the constant mutation rate identified by Drake in his original study of the relationship between mutation rate and genome size[4].

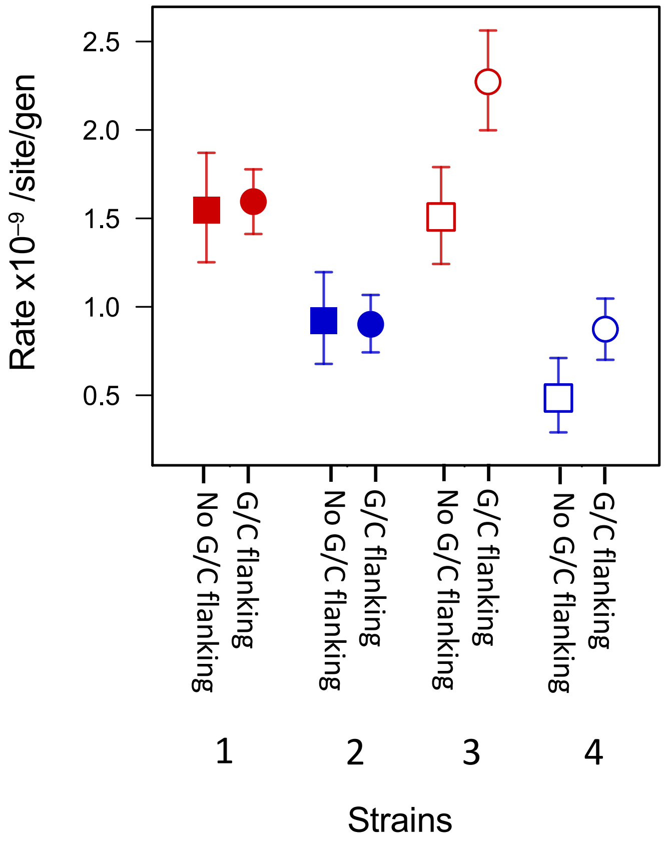


**Fig. S9. The influence of flanking sites on mutation rates for the four strains in the 200-day experiment.** Estimates of mutation rates for sites that have at least one G/C flanking site, and estimates of rates for sites that have no G/C flanking site for each of the four strains from the longer experiment. Points represent mean values across 50 lines, and bars represent 95% confidence intervals from bootstrapping across lines.

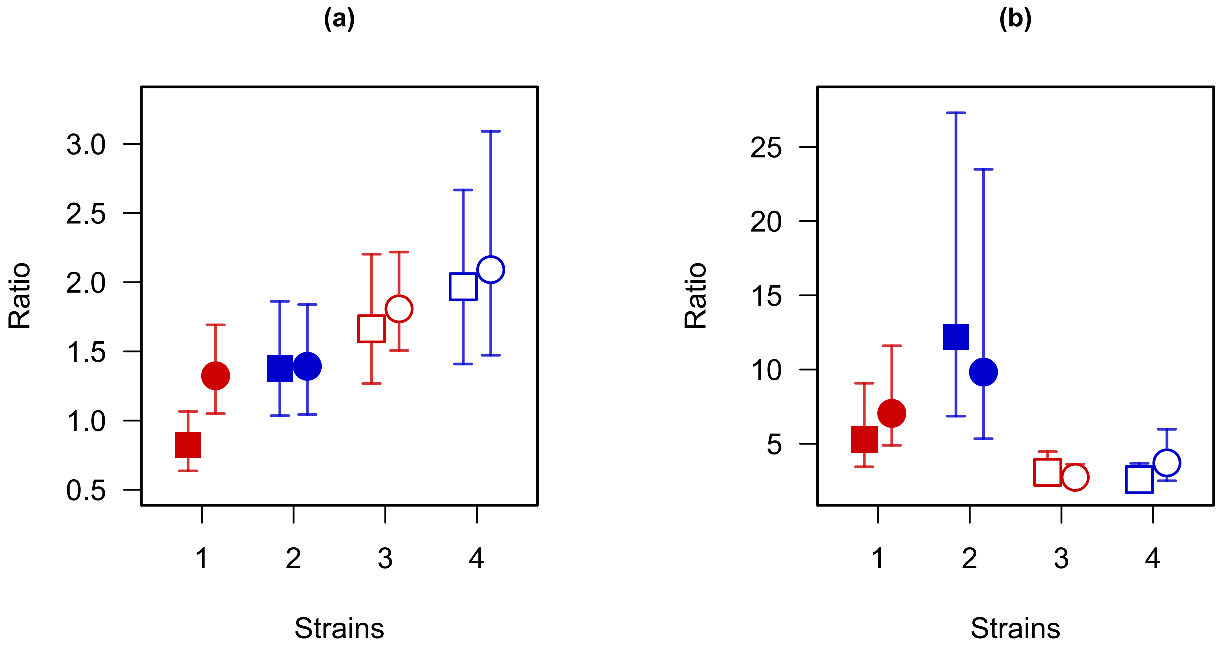


**Fig. S10. Mutational biases in shared and non-shared genomic regions in the four strains in the 200-day experiment.** (a) The ratio of transitions to transversions for shared (circles) and non-shared (squares) regions of the genome. (b) The ratio of G/C to AT transitions to A/T to G/C transitions for non-shared (squares) and shared (circles) regions of the genome. Disease-associated strains are represented by red circles and carriage strains as blue squares. All points represent mean values across 50 lines, and bars represent 95% confidence intervals estimated by bootstrapping.

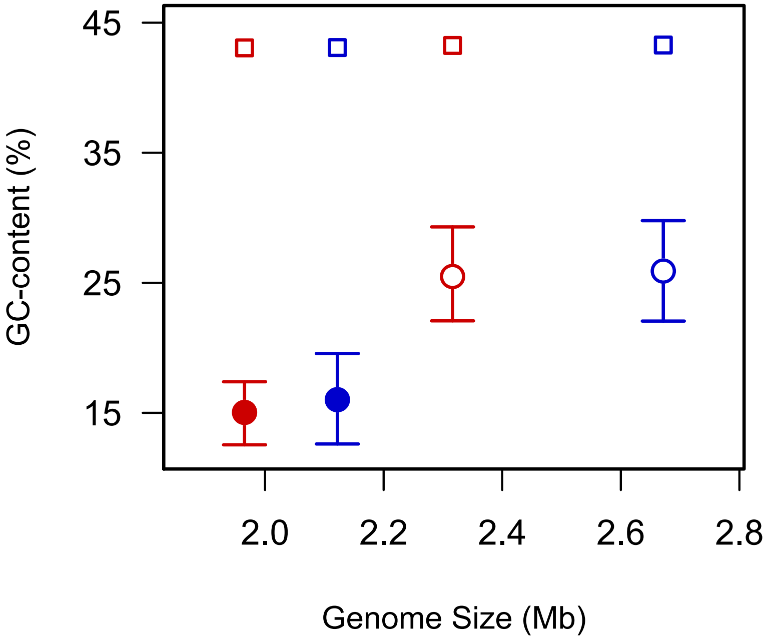


**Fig. S11. Equilibrium GC-content for the four strains in the 200-day MA experiment.** Equilibrium GC-content estimates, calculated from the rates of A/T to G/C mutation and G/C to A/T mutation (circles) and actual genome-wide GC-content for the four strains (squares). Disease-associated strains are represented by red circles and carriage strains as blue squares. All points represent mean values across 50 lines, and bars represent 95% confidence intervals estimated by bootstrapping.

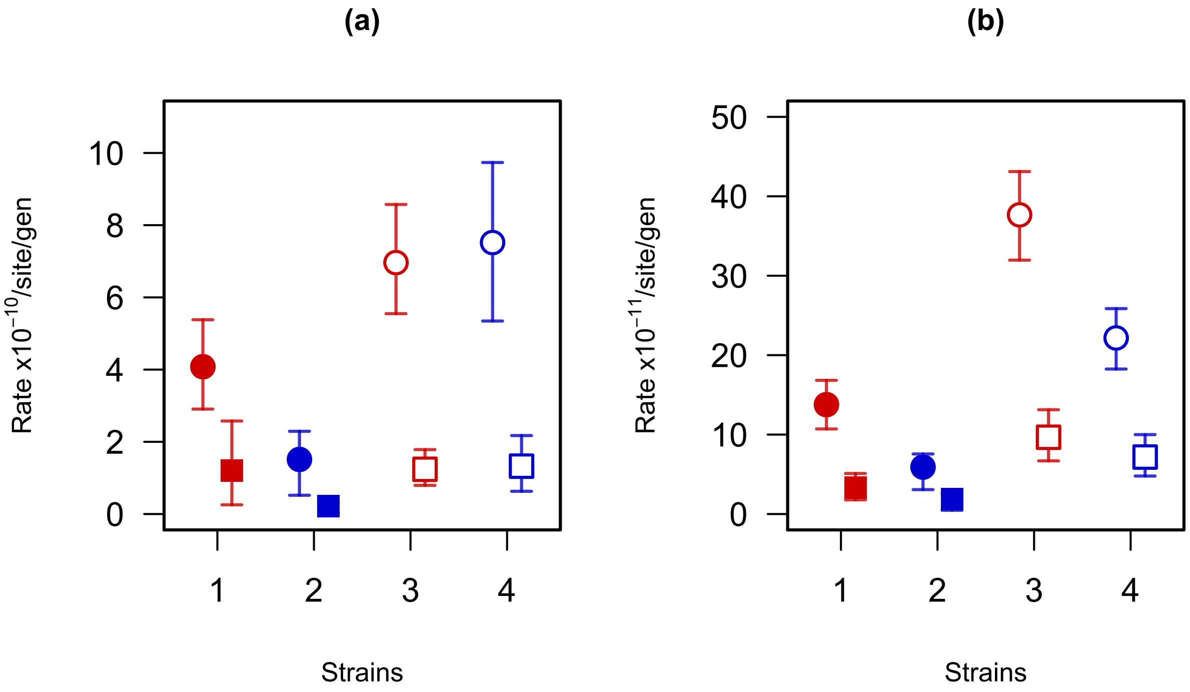


**Fig. S12. Rates of accumulation of small indels for the 200-day MA experiment.** (a) The rates of loss/gain of nucleotide bases through short deletion (circles) and insertion (square) events, and (b) the rates of short deletion (circle) and insertion (squares) events. All points represent mean values across 50 lines, and bars represent 95% confidence intervals estimated by bootstrapping.

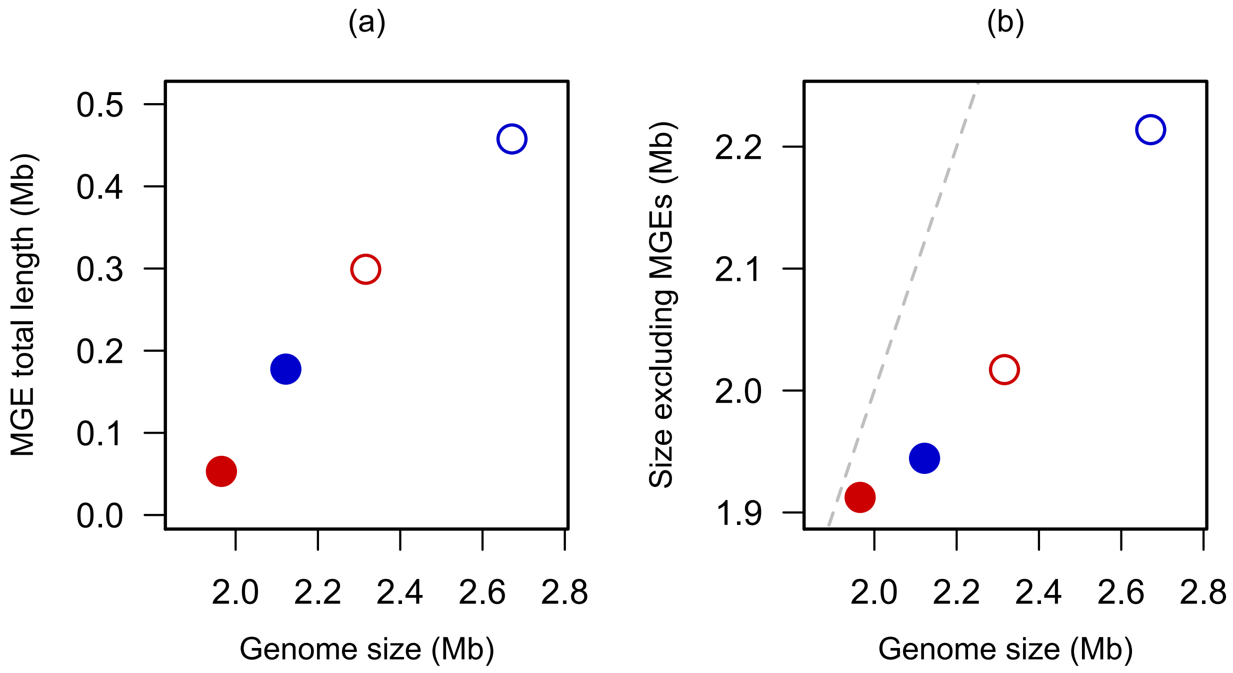


**Fig. S13. Length of elements identified as mobile genomic islands by IslandViewer for the four strains in the 200-day MA experiment.** Figures show the relationship between the total length of mobile genetic elements in each strain and total genome size (a), and the relationship between the length of the genome excluding mobile genetic elements and total genome size (b).

**Table S1. Description of the eight strains used in both MA experiments.**

| **Disease-association** | **Disease** | **Carriage** | **Disease** | **Carriage** | **Disease** | **Carriage** | **Disease** | **Carriage** |
|---|---|---|---|---|---|---|---|---|
| **Genetic background** | **More pathogenic clade** | | | | | | **Less pathogenic clade** | |
| Strain | 1 | 5 | 6 | 2 | 7 | 8 | 3 | 4 |
| Country of origin | Canada | Spain | Netherlands | Canada | Denmark | Canada | Canada | UK |
| Year of sampling | 2014 | 2015 | 2017 | 2016 | 2018 | 2016 | 2009 | 2013 |
| Cluster (1) | 1 | | 12 | | 3 | | 10 | |
| No. lines sequenced | 50 | 13 | 12 | 50 | 13 | 11 | 50 | 50 |
| Genome size (Mb) | 1.97 | 2.00 | 2.11 | 2.12 | 2.22 | 2.10 | 2.32 | 2.67 |
| No. genes | 1952 | 1964 | 2055 | 2120 | 2241 | 2144 | 2344 | 2712 |
| Length coding (Mb) | 1.74 | 1.76 | 1.83 | 1.89 | 1.96 | 1.90 | 2.09 | 2.36 |
| %GC: core genome | 0.4309 | 0.4309 | 0.4309 | 0.4310 | 0.4311 | 0.4312 | 0.4330 | 0.4330 |
| Generation time (mins) | 70 | 67 | 68 | 72 | 67 | 68 | 81 | 80 |
| Previous strain names and references | 1619952 ref.[5] | - | - | DB1V3-4A ref.[5] | - | MZ1B3-4E ref.[5] | 1191316 ref.[5] | 684-21B ref.[6] |
| Long-read data | ENA: SAMEA5610089 (PRJEB21775) | MicrobesNG: 5304491C-278D-4B29-AA2A-D67A13DEF6B5 | MicrobesNG: 5304491C-278D-4B29-AA2A-D67A13DEF6B5 | ENA: SAMEA5610090 (PRJEB21775) | MicrobesNG: 08410916-9F24-47E7-9412-5AC9E3ADAD7B | MicrobesNG: 08410916-9F24-47E7-9412-5AC9E3ADAD7B | ENA: SAMEA5610088 (PRJEB21775) | ENA: SAMEA5610091 (PRJEB21775) |
| Short-read data | ref. [5] | MicrobesNG: 5304491C-278D-4B29-AA2A-D67A13DEF6B5 | MicrobesNG: 5304491C-278D-4B29-AA2A-D67A13DEF6B5 | ref. [5] | MicrobesNG: 08410916-9F24-47E7-9412-5AC9E3ADAD7B | MicrobesNG: 08410916-9F24-47E7-9412-5AC9E3ADAD7B | ref. [5] | ref. [6] |

**Table S2 (separate file). Counts of colony forming units (CFUs) after 24 hours of growth for the 8 ancestral strains used to estimate generation time for each strain.**

**Table S3 (separate file). Description of all single-base mutations observed for each of the 50 lines of the 4 strains in the 200-day experiment.** The table describes the position of the mutations in the (single contig) assembly of the ancestral strain, the inferred single base change, the category of site, whether or not the site is shared across strains ('Shared region'), whether the site is on the leading or lagging strand, the two flanking bases, and whether or not the mutation is clustered (within 30 bp of another mutation in that line). The site categories are 1st/2nd codon positions (1), 3rd codon positions (3), 4-fold degenerate sites (4) and intergenic (0). Shared sites (1) are defined by whether or not the gene is present in all strains for sites within genes, and whether or not the two flanking genes are present in all strains for intergenic regions.

**Table S4. Results of Chi-Squared test for the goodness of fit of a Poisson distribution to mutation rates of single-base substitutions in the 200-day MA experiment.** Counts were divided into four categories for each strain, with each category having an expected frequency >5 under a Poisson distribution given our estimates of the mean rate.

| **Strain** | **Single-base substitutions** | | |
|---|---|---|---|
| | $X^2$ | *df* | *p* |
| **1** | **4.21** | **3** | **0.24** |
| **2** | **1.49** | **3** | **0.71** |
| **3** | **1.48** | **3** | **0.69** |
| **4** | **6.10** | **3** | **0.11** |

**Table S5 (separate file). Growth rates for ancestral and evolved lines in the 200-day MA experiment.**

**Table S6. Comparison of numbers of single-base mutations observed at day 100 and day 200 in the 200-day experiment.** 5 lines of each strain in the 200-day MA experiment were sequenced at the mid-point of the experiment to establish whether faster rates were transitory. Lines were selected that had higher than average rates from each strain. We found no evidence of a difference in rate over the first and second half of the experiment.

| | Number of SNPs | |
|---|---|---|
| **Strain** | **200 days** | **100 days** |
| **1** | 26 | 12 |
| | 22 | 12 |
| | 12 | 6 |
| | 17 | 9 |
| | 24 | 14 |
| **2** | 22 | 10 |
| | 16 | 6 |
| | 16 | 9 |
| | 14 | 1 |
| | 10 | 10 |
| **3** | 26 | 10 |
| | 22 | 10 |
| | 21 | 9 |
| | 21 | 10 |
| | 17 | 6 |
| **4** | 21 | 10 |
| | 21 | 13 |
| | 21 | 15 |
| | 20 | 7 |
| | 18 | 5 |

**Table S7. Core genome pairwise nucleotide distances between the eight ancestral strains used in the MA experiments.** The proportion of nucleotide bases that differ between pairs of strains in the MA experiments based on an alignment of shared genes generated by Panaroo. Closely related disease/carriage pairs are highlighted.

| | 1 (disease) | 5 (carriage) | 6 (disease) | 2 (carriage) | 8 (disease) | 7 (carriage) | 3 (disease) | 4 (carriage) |
|---|---|---|---|---|---|---|---|---|
| **1** | | $1.20 \times 10^{-4}$ | $1.78 \times 10^{-2}$ | $1.78 \times 10^{-2}$ | $3.02 \times 10^{-2}$ | $3.03 \times 10^{-2}$ | $5.49 \times 10^{-2}$ | $5.62 \times 10^{-2}$ |
| **5** | | | $1.78 \times 10^{-2}$ | $1.78 \times 10^{-2}$ | $3.02 \times 10^{-2}$ | $3.02 \times 10^{-2}$ | $5.49 \times 10^{-2}$ | $5.62 \times 10^{-2}$ |
| **6** | | | | $7.39 \times 10^{-5}$ | $2.92 \times 10^{-2}$ | $2.91 \times 10^{-2}$ | $5.45 \times 10^{-2}$ | $5.57 \times 10^{-2}$ |
| **2** | | | | | 2.92x10-2 | $2.91 \times 10^{-2}$ | $5.45 \times 10^{-2}$ | $5.57 \times 10^{-2}$ |
| **8** | | | | | | $1.06 \times 10^{-3}$ | $5.53 \times 10^{-2}$ | $5.61 \times 10^{-2}$ |
| **7** | | | | | | | $5.53 \times 10^{-2}$ | $5.62 \times 10^{-2}$ |
| **3** | | | | | | | | $4.85 \times 10^{-2}$ |
| **4** | | | | | | | | |

**Table S8 (separate file). Description of all single-base mutations observed for each of the lines of the 4 strains in the shorter experiment.**

**Table S9 (separate file). Description of all short indel mutations observed for each of the 50 lines of the 4 strains.**

**Table S10 (separate file). Description of all long deletion mutations observed for each of the 50 lines of the four strains.** Each row is a line in which a long deletion was identified. Lines in which no long deletion was identified are not described. The locations and lengths of the deletions are described, and genes in the deleted regions described. Whether or not the deleted region was identified as a mobile genetic element (MGE) by IslandViewer is described (1=yes).

**Table S11. The size of regions identified as mobile genetic elements for each strain in the 200-day MA experiment.** The lengths of the regions identified by IslandViewer for each strain.

| **Strain** | **Number of genes in mobile elements** | **Length of mobile elements, bp (%)** |
|---|---|---|
| Small-genome pathogen | 58 | 53,085 (2.7 %) |
| Small-genome carriage | 205 | 177,561 (8.3 %) |
| Large-genome pathogen | 271 | 299,087 (12.9 %) |
| Large-genome carriage | 505 | 457,681 (17.1 %) |

**Table S12 (separate file). The regions identified as mobile genetic elements by IslandViewer.**

**Table S13 (separate file). Table of gene presence/absence across 8 strains.** The output of Panaroo for the eight strains used in our MA experiments.

**Table S14 (separate file). Genes in major DNA repair pathways in the 8 strains.**

**Table S15. Oligonucleotide primers (Sigma-Aldrich) and conditions used for Multiplex PCR strain identification.**

<table>
<tr><th colspan="6">Primers</th><th rowspan="3">Thermocycling conditions</th><th rowspan="3">PCR reagents per sample (10 µl)</th></tr>
<tr><th rowspan="2">Name</th><th rowspan="2">Sequence (5'-3')</th><th colspan="4">Product size (bp)</th></tr>
<tr><th>Strain 1</th><th>Strain 2</th><th>Strain 3</th><th>Strain 4</th></tr>
<tr><td>MA-F2</td><td>ATACTCAATGAAA ATCAGAATCAGA</td><td>570</td><td>570</td><td>730</td><td>840</td><td rowspan="6">95°C - 10min<br>x1 cycle<br><br>95°C - 30s<br>53°C - 30s<br>72°C - 1min<br>x 30 cycles<br><br>72°C 5min<br>x1 cycle</td><td rowspan="6">1 µl DNA template<br>5.3375 µl MilliQ H2O<br>2 µl MyTaq Red Reaction Buffer<br><br>0.375 µl of the primer<br><br>0.0625 µl MyTaq DNA polymerase<br><br>0.1 µl DMSO</td></tr>
<tr><td>MA-R2</td><td>TCGGTCAAAT GGTTTGTCG</td><td>570</td><td>570</td><td>730</td><td>840</td></tr>
<tr><td>MA-F5</td><td>ATTTGATGACT GAAAAGCTCCT</td><td>-</td><td>234</td><td>284</td><td>-</td></tr>
<tr><td>MA-R5</td><td>GGTTTGAAGTT ATAATAAAAAACAA</td><td>284</td><td>234</td><td>284</td><td>234</td></tr>
<tr><td>SsuisF</td><td>CTGTAAACCAAT CCATCTTGA</td><td>853</td><td>853</td><td>853</td><td>853</td></tr>
<tr><td>SsuisR</td><td>CTCATTTCAAGG GCAGATAC</td><td>853</td><td>853</td><td>853</td><td>853</td></tr>
</table>

**Table S16 (separate file). The relationship between CFU count and OD during exponential growth in the four ancestral strains from the 200-day experiment**. The relationship between OD and CFU count can vary across bacterial strains. To test whether OD is a reliable indicator of CFU count in the strains in our 200-day experiment, we undertook an additional growth rate experiment following the same procedure as before. Every half hour for the first 5 hours of growth, then every hour up until 8 hours of growth, and at the end of the experiment (24 hours of growth), 100µl of the 300µl culture was removed from a single well. This was serially diluted and spot plated on three THY plates. Comparisons of OD and CFU count revealed a linear relationship in all four ancestral strains during the period of exponential growth (between 0.5 and 3.5 hours). For all four strains OD was found to be a good predictor of change in CFU count during exponential growth (Pearson's correlation coefficient >0.9 for each strain).

### References


1. Murray, G. G. R. *et al.* Genome reduction is associated with bacterial pathogenicity across different scales of temporal and ecological divergence. *Molecular Biology and Evolution* **38**, 1570–1579 (2021).
2. Lynch, M. *et al.* Genetic drift, selection and the evolution of the mutation rate. *Nature Reviews Genetics* vol. 17 704–714 (2016).
3. Senra, M. V. X. *et al.* An unbiased genome-wide view of the mutation rate and spectrum of the endosymbiotic bacterium *Teredinibacter turnerae*. *Genome Biology and Evolution* **10**, 723–730 (2018).
4. Drake, J. W. A constant rate of spontaneous mutation in DNA-based microbes. *Proceedings of the National Academy of Sciences* **88**, 7160–7164 (1991).
5. Hadjirin, N. F. *et al.* Linking phenotype, genotype and ecology: antimicrobial resistance in the zoonotic pathogen *Streptococcus suis*. *bioRxiv* 2020.05.05.078493 (2020) doi:10.1101/2020.05.05.078493.
6. Wileman, T. M. *et al.* Pathotyping the Zoonotic Pathogen *Streptococcus suis*: Novel genetic markers to differentiate invasive disease-associated isolates from non-disease-associated isolates from England and Wales. *Journal of Clinical Microbiology* **57**, (2019).
